# Large generative mRNA language foundation model for efficient coding sequence generation and design with mRNA-GPT

**DOI:** 10.64898/2025.12.22.695962

**Authors:** Bian Bian, Yiming Zhang, Hongmin Li, Jiuzhou Zhong, Yutaka Saito

## Abstract

mRNA design plays a central role in synthetic biology, nucleic acid therapeutics, and vaccine development. Although large language models are applied in many biological fields, generative language models for *de novo* mRNA design remains largely unexplored. Here, we introduce mRNA-GPT, a series of generative mRNA language models which for the first time covers the three domains of life as pretraining datasets. Based on a GPT-2 transformer architecture with 302 million parameters, we pre-trained three separate models on 19,676 bacterial, 4,688 eukaryotic, and 702 archaeal species, leveraging 80 million, 83 million, and 2 million mRNA coding sequences, respectively. Distinct clustering of mRNA coding sequence embeddings from animals, plants, and fungi in the pretrained mRNA-GPT-eukaryote indicates that the model captures organism-specific sequence features. Following unsupervised pre-training, we fine-tuned mRNA-GPT on a translation-efficiency dataset to generate high-performance mRNA sequence. Compared to the pretrained model, the fine-tuned mRNA-GPT produced mRNA sequences with significantly higher translation efficiency scores, demonstrating the ability of mRNA-GPT to capture sequence features underlying high translation efficiency. We further fine-tuned mRNA-GPT on datasets for mRNA stability and mRNA expression, where it likewise produced high-performance mRNA sequences. Our pretrained models are publicly available, enabling other researchers to adapt mRNA-GPT to specialized tasks such as tissue-specific mRNA expression or stability by fine-tuning on their own data. Together, our study demonstrates that generative mRNA language modeling as a promising approach for accelerating mRNA design across diverse biological fields.

## Introduction

Messenger RNA (mRNA) design is a cornerstone of synthetic biology, nucleic acid therapeutics, and vaccine development, and thus plays a critical role in modern biotechnology^1–3^. However, effective mRNA design requires balancing multiple, often competing constraints, including codon usage patterns, RNA secondary structure, RNA modification landscapes, and host-specific regulatory mechanisms^1,4,5^. Achieving mRNA sequences that are both highly expressible and stable within the intended biological context remains a challenging task^1,6^. Although billions of mRNA sequences are now available across the tree of life, transforming this vast corpus into actionable design principles is still difficult, as identifying suitable mRNA sequences require an enormous search space^4,7^.

Large language models (LLMs) have demonstrated outstanding performance in natural language processing (NLP), showing exceptional capability and flexibility in managing complex linguistic tasks^8–10^. Given the power of LLMs, recently they have begun to be applied to biological sequences^11^. These LLMs have transformed representation learning in biology by extracting rich, context-aware features directly from raw sequences without the need for manual annotation^12,13^. Recently, large-scale protein language models have been developed, which are pretrained on millions of unlabeled protein sequences to learn universal embeddings and subsequently fine-tuned for various downstream protein tasks^14–16^. Representative examples include the ESM series and ProteinBERT, which have advanced protein sequence understanding and structure prediction by leveraging BERT^14,17,18^. Furthermore, recent advances in generative protein language models including autoregressive frameworks such as ProtGPT2 for *de novo* protein design and novel Hyena-based architectures like ProtHyena have significantly enhanced the scalability and efficiency of modeling long protein sequences^19,20^.

In the field of nucleotide language modeling, the DNA language models such as the DNABERT series employ a bidirectional encoder architecture to capture contextual patterns within nucleotide sequences and facilitate genomic sequence prediction^13,21^. Recently, generative DNA language models such as Evo and Evo2 have been developed for the design of functional biological systems and *de novo* genome generation, further demonstrating the potential of sequence-based generative modeling^22,23^. Messenger RNA (mRNA) plays a central role in the flow of genetic information from DNA to protein^24^. However, most RNA-focused language models have concentrated on non-coding RNA (ncRNA)^25^. Models such as RNA-FM, RNAErnie, RNABERT and LncRNA-BERT, based on the BERT architecture and pretrained on large ncRNA datasets, have been primarily used for RNA secondary/tertiary structure prediction and protein-RNA interaction prediction and long ncRNA classification^26–29^. In addition, the generative RNA language model GenerRNA which pre-trained on approximately 16 million ncRNA sequences, has been applied to *de novo* RNA design with stable secondary structures and can be fine-tuned to generate novel RNA sequences exhibiting high binding affinity to target proteins^30^. However, mRNA-specific language models remain relatively underexplored. Recent studies, such as CodonBERT and Plant-FM, have employed the BERT framework to enhance mRNA sequence understanding within a limited number of organisms and to identify functional RNA sequence and structural motifs in plants, respectively^31,32^. Recently, codonGPT was developed using protein-coding nucleotide sequences from 7 model organisms^33^. Nevertheless, large-scale generative mRNA language foundation modeling for *de novo* mRNA sequence design and generation across diverse taxa remains largely unexplored.

In this study, we introduced mRNA-GPT, the first generative mRNA language foundation model which adopts a GPT-2 based Transformer generative architecture with 302M parameters and is pretrained in a domain-aware manner on large, curated mRNA coding sequence corpora: 19,676 bacterial species (80M mRNA sequences), 4,688 eukaryotic species (83M sequences), and 702 archaeal species (2M sequences). Training the three domain models separately allows the mRNA-GPT series to encode lineage-specific regularities such as codon usage regimes and compositional preferences. Beyond large-scale pretraining, mRNA-GPT supports task-conditioned mRNA generation via lightweight fine-tuning. In this setting, property-relevant datasets (e.g., translation efficiency, mRNA stability, or expression) provide supervision that steers sequence generation toward user-specified objectives while retaining naturalness and organism specificity. The resulting framework is designed to accommodate diverse downstream use cases, from host-optimized coding sequences to application-specific constraints in mRNA therapeutics and vaccines.

## Results

### Architecture of mRNA-GPT and the framework for mRNA coding sequence design

mRNA design typically involves both the optimization of untranslated regions (UTRs) and coding sequences (CDSs)^1^. While many well-established commercial UTR sequences are available, optimizing the coding region remains challenging due to codon degeneracy, the phenomenon where multiple codons encode the same amino acid, resulting in an enormous combinatorial search space for identifying the optimal mRNA coding sequence^1^(Figure 1A). Optimization of codon usage patterns is critical for improving translation efficiency, stability, and expression performance in synthetic mRNA design. To address this issue, we adopt a “learning from nature” approach by leveraging large generative language model trained on naturally occurring mRNA sequences to guide *de novo* mRNA coding sequence design. In this study, we developed mRNA-GPT, a GPT-2 based autoregressive large language model trained on large-scale mRNA coding sequences collected from the NCBI RefSeq database, encompassing eukaryotic, bacterial, and archaeal species. As illustrated in Figure 1B, raw mRNA sequences from each biological domain were tokenized at the codon level and used to pretrain the GPT-2 backbone with 302 million parameters, which consists of standard transformer blocks incorporating masked multi-head attention, layer normalization, feed-forward networks, and positional embeddings. This architecture enables the model to capture long-range contextual dependencies within mRNA sequences, facilitating the generation of biologically plausible and highly expressible synthetic mRNAs. To capture domain-specific mRNA coding patterns, we independently pretrained three models for each biological domain, referred to as mRNA-GPT-bacteria, mRNA-GPT-eukaryote, and mRNA-GPT-archaea. The reference mRNA coding sequences were collected from 19,676 bacterial, 4,688 eukaryotic, and 702 archaeal species, respectively. In total, approximately 80 million bacterial, 83 million eukaryotic, and 2 million archaeal sequences were used for model pretraining and validation in each domain. (Figure 1B). The training loss and perplexity curves for each pretrained model are shown in Figure S1, Figure S2, and Figure S3, respectively.

**Figure 1.**
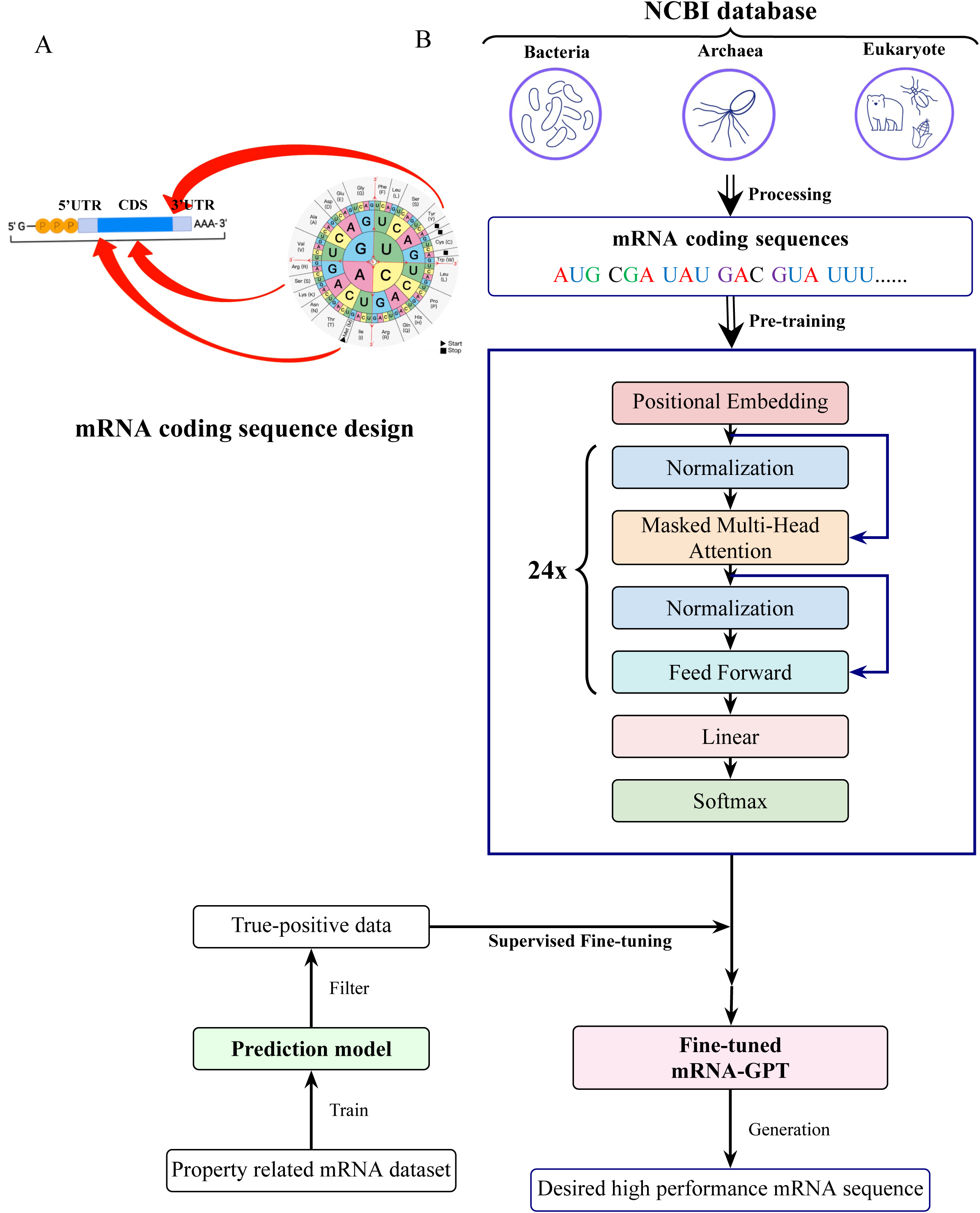
Overview of the mRNA-GPT framework for mRNA coding sequence design. **(A)** Schematic illustration of the full-length mRNA architecture, consisting of the 5′ untranslated region (5′UTR), coding sequence (CDS), and 3′ untranslated region (3′UTR). The coding sequence design follows the standard genetic code, as illustrated by the codon wheel, where each triplet codon corresponds to a specific amino acid. **(B)** The workflow of mRNA-GPT. mRNA coding sequences from bacteria, archaea, and eukaryotes were collected from the NCBI database and preprocessed for large-scale pretraining. The mRNA-GPT model, built upon a 24-layer transformer architecture, was pretrained to capture universal codon-level representations. In the fine-tuning phase, property-related mRNA datasets were used to train a prediction model that filters true-positive data, which were then employed for supervised fine-tuning of the pretrained model. The fine-tuned mRNA-GPT models subsequently enables *de novo* generation of high-performance mRNA sequences with desired biological properties, supporting end-to-end mRNA coding sequence design.

Following the pretraining, the pretrained model was fine-tuned through supervised fine-tuning (SFT) on curated datasets comprising experimentally validated, high-performance mRNA transcripts, enabling the generation of biologically meaningful and highly expressible sequences. Firstly, property-specific datasets (mRNA translation efficiency, stability, and expression) were used to train a prediction model that filtered high-confidence (“True-Positive”) samples (see Method). These curated data were subsequently employed to fine-tune the pretrained model. After fine-tuning, the generated mRNA sequences were evaluated by the prediction model and the top-ranked candidates were selected as high-property mRNA designs (Figure 1B). This approach established a closed-loop framework that integrates generation, prediction, and optimization, thereby facilitating functional mRNA coding sequence design in an end-to-end manner.

### The pretrained mRNA-GPT learned mRNA coding pattern within three domains of life

To investigate whether mRNA-GPT effectively learns biologically meaningful representations of mRNA coding sequences, we analyzed the latent embeddings obtained from the final hidden states of pretrained models using principal component analysis (PCA) as preprocessing step followed by uniform manifold approximation and projection (UMAP) visualization (Figure 2A). Distinct clustering patterns were observed for each of three biological domains (eukaryotes, bacteria, and archaea).

**Figure 2.**
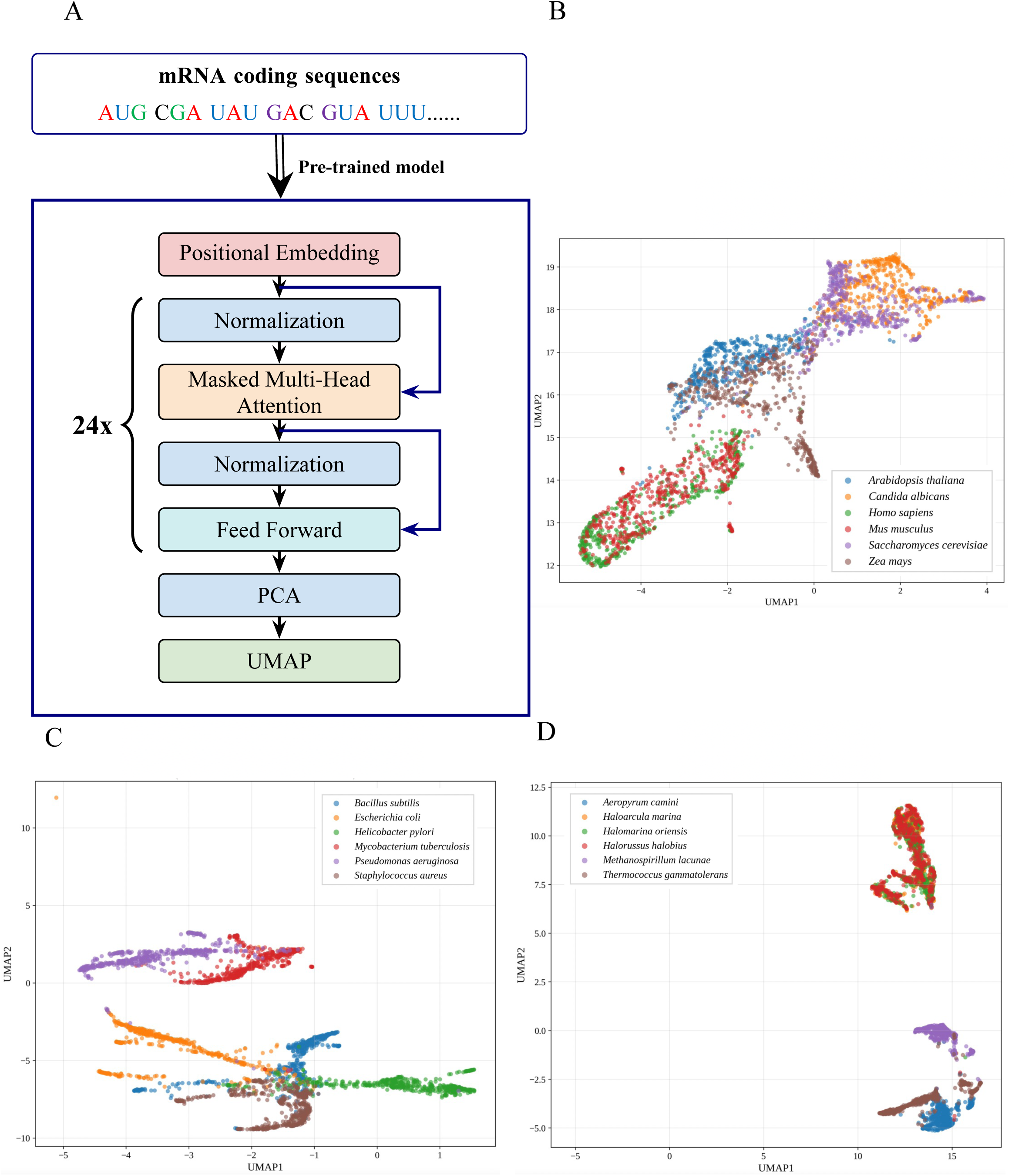
mRNA-GPT leaned mRNA coding patterns across three biological domains. (A) Two-dimensional UMAP plots display RNA sequence embeddings derived from the last hidden states of the pretrained mRNA-GPT models for eukaryotes (B), bacteria (C), and archaea (D). Each point represents an mRNA coding sequence, with its color indicating the species of origin. For each species, 500 mRNA coding sequences were used.

As shown in the UMAP visualization, the clusters of human (green) and mouse (red) are positioned closely together, both belonging to the animal kingdom, indicating that the model successfully captured the high similarity in their mRNA sequence features. Arabidopsis (blue) and maize (gray) are distributed on the opposite side, adjacent to each other but clearly separated from the animal cluster, consistent with their phylogenetic proximity within the plant kingdom. Yeast (purple) and *Candida albicans* (orange) form distinct clusters in the upper region, separated from both plants and animals, reflecting the unique mRNA characteristics of fungi. This distribution pattern is highly consistent with the phylogenetic tree of eukaryotes, suggesting that the latent representations learned by mRNA-GPT implicitly encode evolutionary relationships and species-specific features (Figure 2B). In bacteria domain, *Escherichia coli* (orange) and *Pseudomonas aeruginosa* (purple) form distinct but nearby clusters, both representing Gram-negative *Proteobacteria* with related sequence features. *Bacillus subtilis* (blue) and *Staphylococcus aureus* (brown), both Gram-positive *Firmicutes*, are positioned closely together in the lower region, reflecting their phylogenetic relatedness and shared codon usage tendencies. *Mycobacterium tuberculosis* (red) lies between the Gram-positive and Gram-negative groups, consistent with its unique high-GC *Actinobacterial* genome. *Helicobacter pylori* (green) appears as a separate cluster at the rightmost region, illustrating its distant evolutionary lineage (*Epsilonproteobacteria*) (Figure 2C). Similarly, in archaea (Figure 2D), closely related halophilic species (*Haloarcula marina*, *Halomarina oriensis*, and *Halorussus halobius*) grouped together, while thermophilic species (*Aeropyrum camini*, *Thermococcus gammatolerans*) occupied distant regions in the embedding space.

Together, these observations demonstrate that mRNA-GPT learns codon-level and compositional patterns reflective of evolutionary and functional similarities, enabling the unsupervised organization of mRNA coding sequences according to taxonomic relationships.

### Fine-tuned mRNA-GPT in bacteria can generate high translation efficiency mRNA sequence

Translation efficiency (TE) plays a critical role in determining protein abundance and expression yield; therefore, improving TE is one of the major objectives in mRNA design^34–36^. As shown in Figure 3A, TE can be quantified using ribosome profiling data.

**Figure 3.**
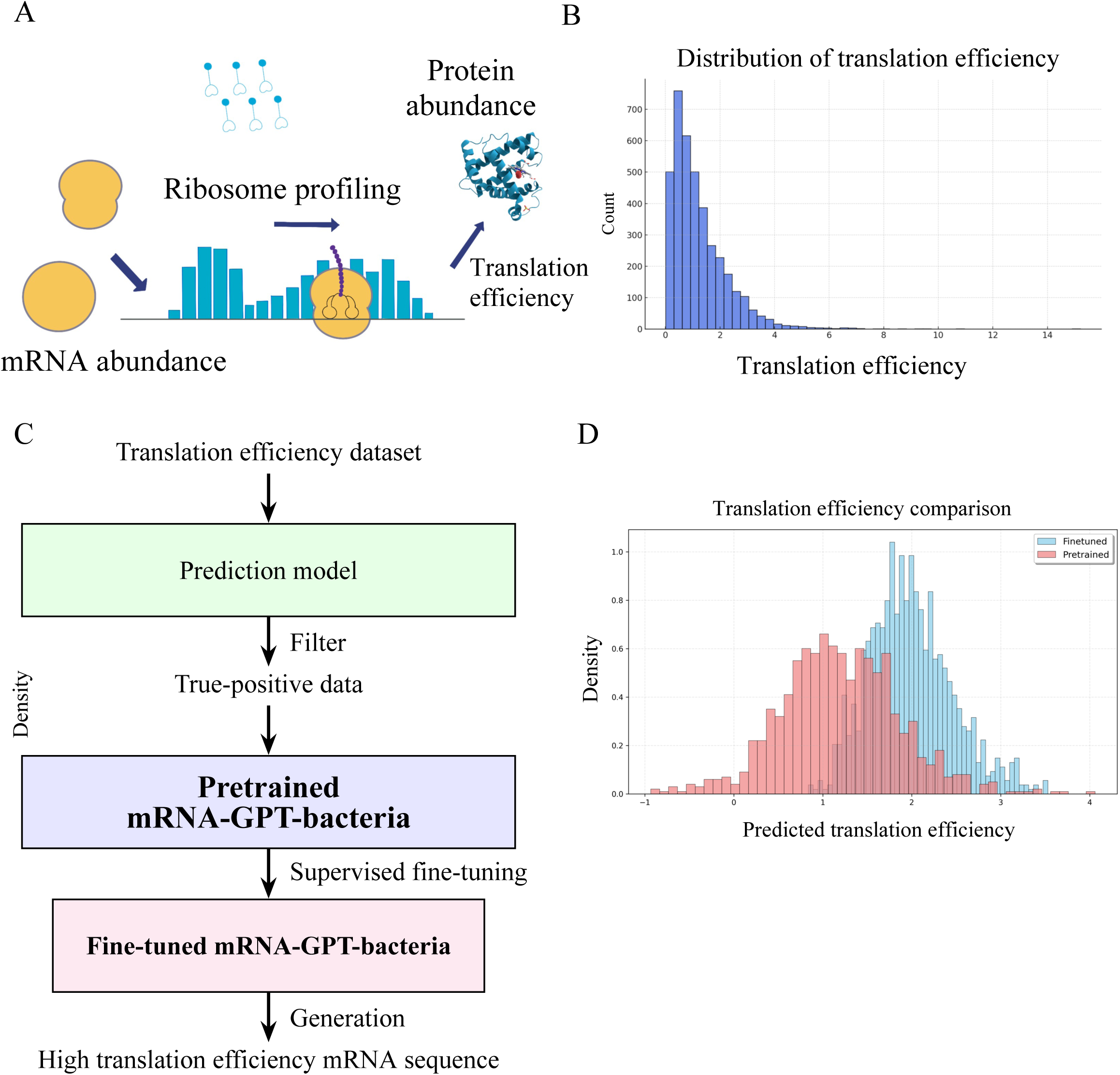
Fine-tuning and evaluation of the mRNA-GPT-bacteria model for mRNA coding sequence design with translation efficiency optimization. (A) Overview of translation efficiency measurement. Translation efficiency (TE) is calculated by integrating mRNA abundance and ribosome profiling data to estimate the relative protein abundance generated per transcript. (B) Distribution of translation efficiency values across *E.coli* genes. (C) Workflow of the mRNA-GPT-bacteria fine-tuning process. A translation efficiency dataset is first used to train a prediction model, which filters true-positive samples for supervised fine-tuning of the pretrained mRNA-GPT-bacteria model. The fine-tuned model is then used for *de novo* generation of mRNA sequences. (D) Comparison of the predicted translation efficiency of mRNA coding sequences generated by the pretrained and fine-tuned models.

To design and generate high TE mRNA coding sequence, firstly, we curated a bacterial translation efficiency dataset (Figure 3B) and examined its sequence length (Figure S4A) and TE value distributions (Figure 3B). We first developed a prediction model named mRNA2TE (Figure S4B,C), which takes mRNA coding sequences as input and predicts their corresponding TE values. Using this model, we inferred the TE for all sequences in the dataset and defined a true-positive subset consisting of sequences whose predicted and experimentally measured TE values were both higher than 2. This high-TE subset was then used for SFT of the pretrained mRNA-GPT-bacteria model (Figure 3C). Given that only 190 sequences met the threshold, we selected the best-performing checkpoint, as further training led to overfitting. To ensure a fair comparison, both pretrained and fine-tuned models were evaluated using datasets with matched sequence length distributions (Figure S7A). As shown in Figure 2D, the TE distribution of the fine-tuned outputs shifts rightward relative to that of the pretrained model, demonstrating that supervised fine-tuning on high-TE samples effectively enables the generation of mRNA sequences with enhanced translational efficiency.

### Fine-tuned mRNA-GPT in archaea generates highly stable mRNA sequences

mRNA stability is another critical factor influencing protein production and yield^37,38^. In this study, we sought to leverage the rich informational resources embedded in archaeal genomes, as archaea thrive in extreme environments such as high temperature and high salinity, which may have driven the evolution of distinctive coding patterns that contribute to mRNA transcript stability^39–41^. These unique sequence characteristics make archaeal genomic data a valuable resource for designing highly stable mRNA sequences (Figure 4A). Therefore, we selected the pretrained mRNA-GPT-archaea model for further fine-tuning.

**Figure 4.**
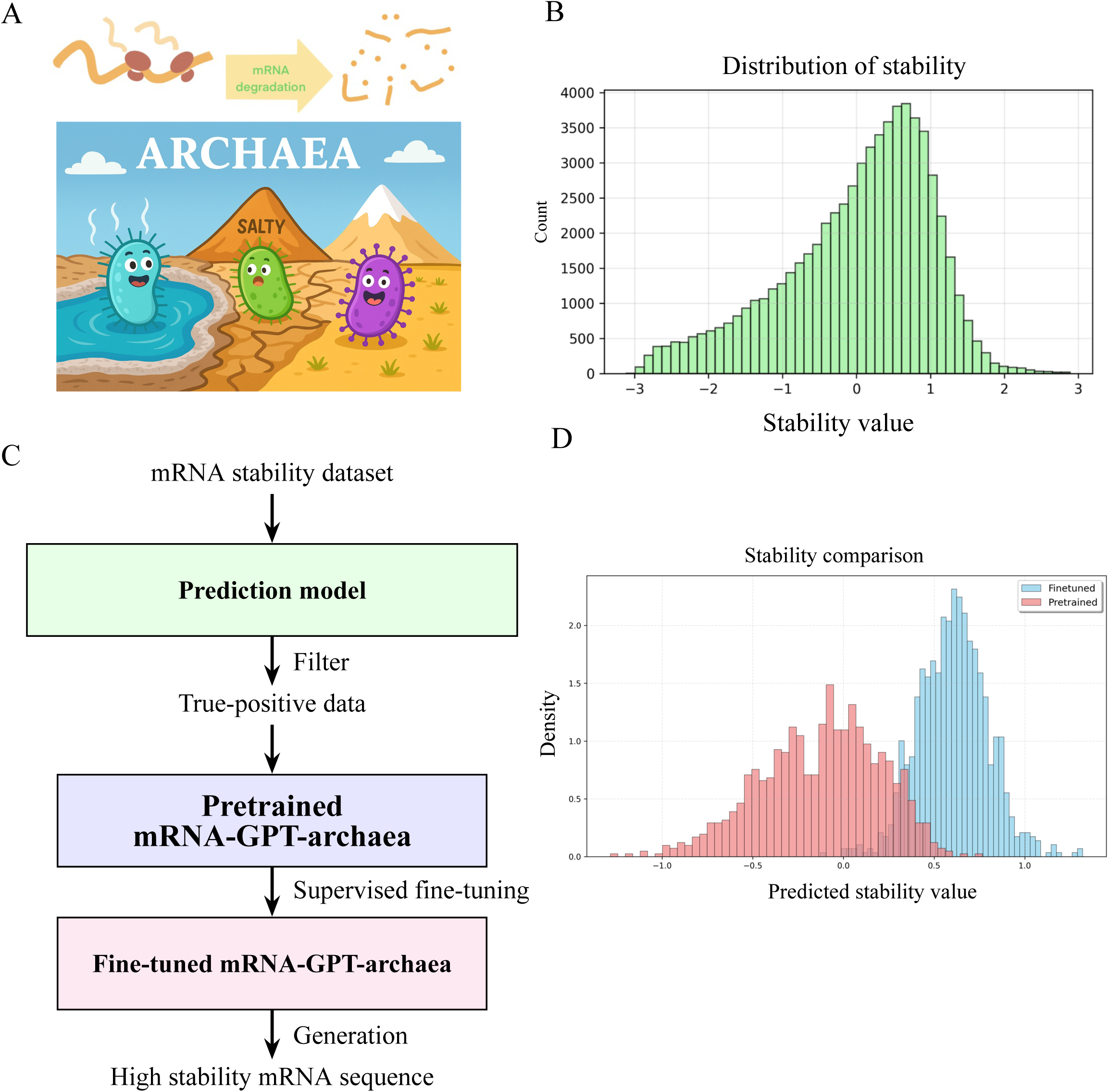
Fine-tuning and evaluation of the mRNA-GPT-archaea model for mRNA coding sequence design with stability optimization. (A) Overview of mRNA degradation and the extreme environments of archaea, illustrating how archaeal genetic information can be leveraged to design high-stability mRNA sequences, the cartoon is made by ChatGPT. (B) Distribution of mRNA stability dataset. (C) Workflow of the mRNA-GPT-archaea fine-tuning process. A mRNA stability dataset is firstly used to train a prediction model, which filters true-positive samples for supervised fine-tuning of the pretrained mRNA-GPT-archaea model. The fine-tuned model is then used for *de novo* generation of mRNA coding sequences. (D) Comparison of the predicted stability value of mRNA coding sequences generated by the pretrained and fine-tuned models.

We firstly selected the mRNA stability dataset (Figure S5A) and examined the distribution of mRNA stability value (Figure 4B). The fine-tuning procedure followed a similar workflow to that used for translation efficiency dataset (Figure 4C). We first developed a prediction model, mRNA2Stability, which takes mRNA coding sequences as input and outputs a predicted stability value (Figure S5B,C). Using this model, we inferred the stability for all sequences in the dataset and defined a true-positive subset consisting of sequences whose predicted and experimentally measured stability values were both higher than 0.5. In this case, approximately 3,000 sequences were used to fine-tune the pretrained mRNA-GPT-archaea model for 30 epochs, the best-performing checkpoint was then selected for downstream sequence generation. To ensure a fair comparison, both pretrained and fine-tuned models were evaluated using datasets with matched sequence length distributions (Figure S7 B). As shown in Figure 4D, the stability distribution of fine-tuned outputs shifted markedly rightward compared with that of the pretrained model, indicating that fine-tuning with high-stability samples effectively guided mRNA-GPT-archaea to generate mRNA sequences with substantially enhanced stability.

### Fine-tuned mRNA-GPT in eukaryote generates high expression mRNA sequences

mRNA expression levels are strongly correlated with protein abundance, as transcripts with higher expression generally yield higher amounts of protein products^38,42^. To explore whether mRNA-GPT can be adapted to design mRNA with enhance expression, we selected a fungal expression dataset as the training corpus, given that fungi are widely used as expression hosts in modern cell factory systems^43,44^.

We first examined the distribution of expression values across the dataset (Figure S6A; Figure 5A) and then trained a prediction model, mRNA2Expression, which predicts expression value directly from mRNA sequences (Figure S6B,C). Using this model, we inferred the expression for all sequences in the dataset and defined a true-positive subset consisting of sequences whose predicted and experimentally measured stability values were both higher than 0.5. The fine-tuning procedure followed a similar workflow to that used for translation efficiency dataset (Figure 5B). About 650 sequences were used to perform SFT of the pretrained mRNA-GPT-eukaryote model and the optimal checkpoint was selected for downstream generation of sequences with enhanced expression potential.

**Figure 5.**
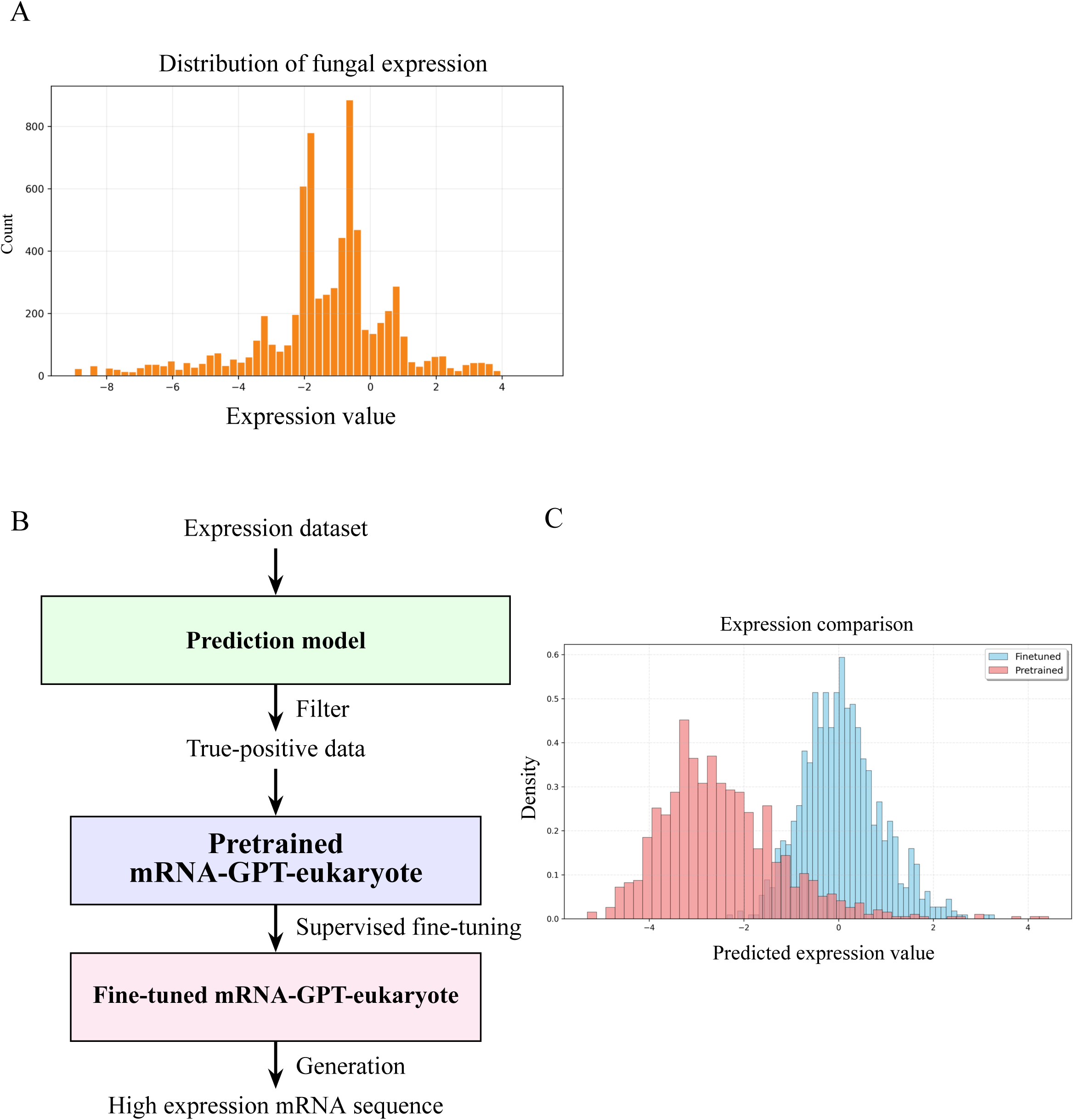
Fine-tuning and evaluation of the mRNA-GPT-eukaryote model for mRNA design with expression optimization. (A) Distribution of mRNA expression dataset. (B) Workflow of the mRNA-GPT-eukaryote fine-tuning process. A mRNA expression dataset is firstly used to train a prediction model, which filters true-positive samples for supervised fine-tuning of the pretrained mRNA-GPT-eukaryote model. The fine-tuned model is then used for *de novo* generation of mRNA coding sequences. (C) Comparison of the predicted expression value of mRNA coding sequences generated by the pretrained and fine-tuned models.

To ensure a fair comparison, both pretrained and fine-tuned models were evaluated using datasets with matched sequence length distributions (Figure S7C). As shown in Figure 5C, the expression value distribution of sequences generated by the fine-tuned model shifts significantly rightward relative to that of the pretrained model, indicating that fine-tuning with high-expression samples effectively guides mRNA-GPT-eukaryote to generate mRNA sequences with markedly enhanced expression capacity.

## Discussion

mRNA design has recently emerged as an active research area, given its tremendous potential in diverse biomedical applications such as influenza vaccines, cancer immunotherapy, and protein replacement therapies^45,46^. Current mRNA design strategies largely depend on algorithmic optimization and conventional rule-based approaches^4,7,47^. However, *de novo* mRNA design driven by generative large language models remains largely unexplored, representing a promising new direction for computational mRNA engineering. In this study, we present mRNA-GPT, the first generative mRNA language model for coding sequence design across the three domains of life including bacteria, archaea, and eukaryotes. Our work demonstrates that generative large language model architectures, when appropriately pre-trained and fine-tuned, can effectively capture biological sequence semantics and generate functional mRNA sequences with enhanced properties such as translation efficiency, stability, and expression. Our framework provides a scalable paradigm for programmable mRNA design, moving beyond traditional rule-based optimization methods.

The pretraining stage of mRNA-GPT plays a central role in enabling the generation of biologically meaningful sequences. The pretraining process of a generative language model is analogous to deeply learning the “dictionary” of a language, acquiring its grammar, syntax, and vocabulary. In contrast, the fine-tuning process corresponds to composing a poem or writing a novel based on that linguistic foundation. Similarly, in the context of the mRNA language, pretraining allows the model to learn the “biological dictionary” embedded in natural mRNA sequences, while fine-tuning enables it to generate new sequences tailored to specific design objectives using the learned biological grammar. The adoption of codon-level tokenization strategy enables the model to preserve the natural compositional units of genetic information, while pretraining on billions of codons from tens of thousands of species allows mRNA-GPT to learn the evolutionary principles underlying codon usage. The three domain-specific pretrained models: mRNA-GPT-bacteria, mRNA-GPT-archaea, and mRNA-GPT-eukaryote aimed to learn domain-specific constraints such as GC bias, codon adaptation index (CAI) patterns, and organism-specific translation machinery preferences. UMAP analysis indicates that pretrained mRNA-GPT indeed learned mRNA sequence features reflecting evolutionary proximity (Figure 2).

To generate mRNA sequence with desired properties, we fine-tuned our mRNA-GPT in bacteria, archaea and eukaryote for translation efficiency (TE), mRNA stability and expression task respectively. Fine-tuned mRNA-GPT-bacteria can generate sequence with enhanced translation efficiency than that of pretrained mRNA-GPT-bacteria (Figure 3). Importantly, this high TE sequence generation was achieved using a relatively small fine-tuning dataset (190 sequences), demonstrating that the pretrained backbone already encoded generalizable biological information that can be efficiently adapted with minimal supervision. Such data-efficient adaptation is a key advantage of generative language model-based frameworks in biological sequence design. Stable transcripts persist longer in the cellular environment, thereby sustaining protein production over extended periods^48,49^. Archaea, which inhabit extreme environments such as high temperature and salinity, offer a unique natural reservoir of stability-optimized genetic sequences^50,51^. To leverage this important genetic resource, we used the pretrained mRNA-GPT-archaea, the resulting fine-tuned model generated sequences with markedly higher predicted stability values, as evidenced by a pronounced rightward shift in the distribution (Figure 4). This suggests that the model successfully captured structural and compositional determinants of mRNA transcript stability. Moreover, mRNA-GPT-eukaryote, extends this framework to the optimization of mRNA expression level, which is a multifactorial property influenced by promoter activity, transcript abundance, and cellular regulation^52,53^. We selected fungi as the fine-tuning system because of their widespread use as expression hosts in industrial biotechnology and recombinant protein production^43^. The fine-tuned mRNA-GPT-eukaryote produced sequences whose predicted expression distribution shifted significantly toward higher values (Figure 5). This suggests that, despite the regulatory complexity of eukaryotic systems, generative mRNA language model can still learn and exploit sequence-encoded determinants of expression, such as codon bias, mRNA structure, and compositional preferences associated with high transcriptional activity.

Despite these promising results, several limitations remain. Our present study focuses on single-objective fine-tuning, biological systems often involve trade-offs between competing properties such as high expression versus low immunogenicity. Extending mRNA-GPT to multi-objective or reinforcement-based training schemes could enable more balanced optimization. Furthermore, experimental validation of the generated sequences will be essential to confirm the biological relevance of the model’s predictions and to ultimately close the loop between *in silico* generation and *in vitro* performance. In addition, a codon-constrained generation algorithm for producing mRNA coding sequences tailored to specific protein targets can be further developed in the future. By applying a dynamic masking strategy to our fine-tuned mRNA-GPT models, the sampling space can be restricted to valid synonymous codons at each position, thereby ensuring 100% translation accuracy while preserving the model’s learned codon-usage preferences. This approach could enable the rational design of mRNA coding sequences for proteins of interest with improved translation efficiency, increased stability, and higher expression levels.

## Material and method

### Datasets used for pretraining and fine-tuning

The mRNA coding sequences used for mRNA-GPT pretraining were sourced from the NCBI repository (https://www.ncbi.nlm.nih.gov/datasets/genome/), accessed in August 2024. Reference mRNA sequences representing bacteria, archaea, and eukaryotes, as curated by RefSeq, were retrieved via the NCBI command-line tools (https://www.ncbi.nlm.nih.gov/datasets/docs/v2/command-line-tools/download-and-install/). Altogether, datasets encompassing 19,676 bacterial, 4,688 eukaryotic, and 702 archaeal species were collected. This process yielded approximately 80 million bacterial, 2 million archaeal, and 83 million eukaryotic mRNA coding sequences, respectively. To finetune the pretrained mRNA-GPT model series for generation task, the translation efficiency dataset^54^, the mRNA stability dataset^55^ and fungal expression dataset^56^ were from were collected, respectively.

### Model architecture of mRNA-GPT

The mRNA-GPT model is constructed using a decoder-only Transformer architecture, adhering to the fundamental design principles of the Generative Pre-trained Transformer (GPT) family^10,57^. Operating as an autoregressive sequence model, it is designed to predict subsequent tokens based on preceding context, effectively capturing the syntactic and semantic dependencies within mRNA coding sequences. The architecture is instantiated with a model dimensionality comparable to GPT-2 Medium, utilizing a hidden size of 1,024 and containing 24 stacked layers^8^.

The processing pipeline begins with the input stage, where discrete sequence data is transformed into continuous representations through two parallel embedding layers. A token embedding layer maps the specialized vocabulary of 69 items into high-dimensional vectors. This vocabulary is specifically constructed to represent the fundamental units of mRNA translation, comprising the five special control tokens [PAD], [UNK], [CLS], [SEP], and [MASK], alongside the complete set of 64 mRNA codons (permutations of A, C, G, and U, such as AAA, AUG, and UUU). Learnable positional embeddings are added to these token vectors to encode the sequential order, allowing the model to handle inputs with a maximum context window of 1,024 tokens and ensuring that long-range dependencies within the mRNA strands are preserved.

The core of the network consists of 24 identical Transformer blocks that employ a pre-normalization strategy, where Layer Normalization is applied to the input of the sub-modules rather than the output to improve training stability^58^. Each block contains a multi-head causal self-attention mechanism with 16 heads, which uses a masking strategy to prevent the model from attending to future tokens. This is followed by a position-wise feed-forward network (MLP) that utilizes a GELU activation function and expands the internal dimension to four times the embedding size. Residual connections are applied across both the attention and MLP layers to facilitate gradient flow through the deep network.

Finally, the output from the last Transformer block is normalized and passed through a linear head to project the hidden states back to the vocabulary dimension for prediction. To maximize parameter efficiency, the weights of this output head are tied to the input token embeddings. The specific configuration of mRNA-GPT optimizes for computational throughput by utilizing Flash Attention and strictly disabling bias terms in all linear and layer normalization modules, resulting in a streamlined and efficient architecture^59,60^.

### Implementation of pretraining mRNA-GPT

We constructed the training corpus through a multi-stage data processing pipeline designed to convert raw genomic information into a clean, codon-based format suitable for autoregressive modeling. Initially, we traversed raw FASTA files to extract nucleotide sequences, applying a rigorous filtering process to ensure biological validity. We retained only those sequences composed exclusively of standard RNA bases (A, U, C, G) and enforced a structural constraint requiring the total nucleotide count to be divisible by three, ensuring alignment with canonical codon triplets. During this transcription phase, we systematically converted thymine (T) residues to uracil (U) to reflect mRNA chemistry and formatted the resulting sequences as space-separated codons.

Following the initial cleaning, we tokenized and serialized the sequences for efficient ingestion. Using a custom tokenizer initialized with our specialized 69-token vocabulary, we converted the codon sequences into integer identifiers. To explicitly model the start and end of translation events, we appended separator tokens ([SEP]) to both extremities of each sequence. We subsequently partitioned the dataset into training and validation subsets using a 90/10 split ratio.

We pretrained the model from scratch for a total of 1,000,000 iterations utilizing a causal language modeling objective, where the network minimizes the cross-entropy loss by predicting the next token in the sequence based on the preceding context. We performed optimization using the AdamW optimizer^61^ (β_1_=0.9, β_2_=0.999) with a weight decay of 0.0, while applying gradient clipping with a threshold of 1.0 to prevent instability. The learning rate followed a cosine decay schedule, initiating with a linear warmup of 5,000 steps to reach a maximum value of 1.0×10^-3^ before decaying to a minimum of 1.0×10^-4^. To handle the computational demands of the dense Transformer architecture, we configured a mini-batch size of 16 with 16 gradient accumulation steps and employed Distributed Data Parallel (DDP) strategies alongside Automatic Mixed Precision (AMP), facilitating efficient scaling across multiple GPU nodes while maintaining numerical precision.

### Fine-tuning process of mRNA-GPT

To enable property-specific mRNA coding sequence optimization, we developed a SFT framework built upon the pretrained mRNA-GPT model. The process consists of three major steps: (1) training a prediction model, (2) filtering high-quality data, and (3) fine-tuning the pretrained generative model. Firstly, a property-related mRNA dataset (e.g., translation efficiency, stability, or expression) was collected and used to train a prediction model capable of estimating the quantitative property score for each mRNA coding sequence. Second, the trained predictor was applied to the entire dataset to evaluate each sequence. Samples exhibiting both high predicted and experimentally measured property values were retained as true-positive data. Specifically, we set the threshold to 2.0 for translation efficiency (TE) and 0.5 for both stability and expression, ensuring that only biologically meaningful and high-quality sequences were included in the subsequent fine-tuning process. Finally, these curated high-confidence sequences were used to perform supervised fine-tuning of the pretrained mRNA-GPT model. During this phase, the model’s generative capability was adjusted to better capture sequence patterns associated with enhanced functional properties. After fine-tuning, the model was employed for *de novo* generation of mRNA coding sequences exhibiting desired biological performance (e.g., high translation efficiency or improved stability).

## Code availability

The codes used for mRNA-GPT pretraining and finetuning can be found at https://github.com/ZHymLumine/mRNA-GPT/

## Acknowledgements

This work was supported by JST CREST JPMJCR23N1. The computations were partially performed on the NIG supercomputer at ROIS National Institute of Genetics, the ABCI supercomputer at AIST, HOKUSHIN supercomputer at Department of Data Science, Kitasato University and the SQUID supercomputer at Cybermedia Center, Osaka University through the HPCI System Research Project (hp230057, hp240075). We thank Prof. Kiyoshi Asai for fruitful discussion.

---

**Figure S1.**
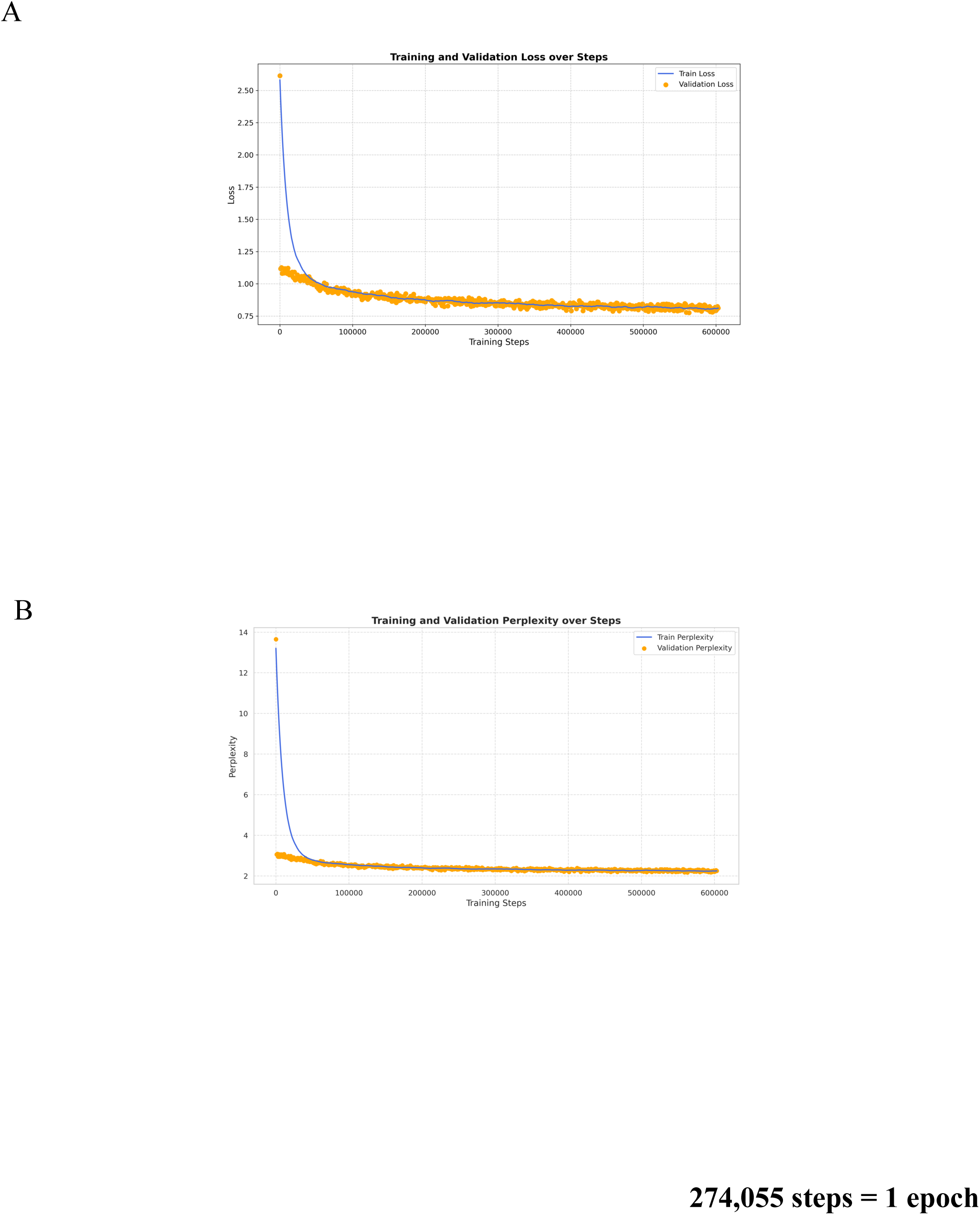
Loss and perplexity during pre-training of mRNA-GPT-bacteria.

**Figure S2.**
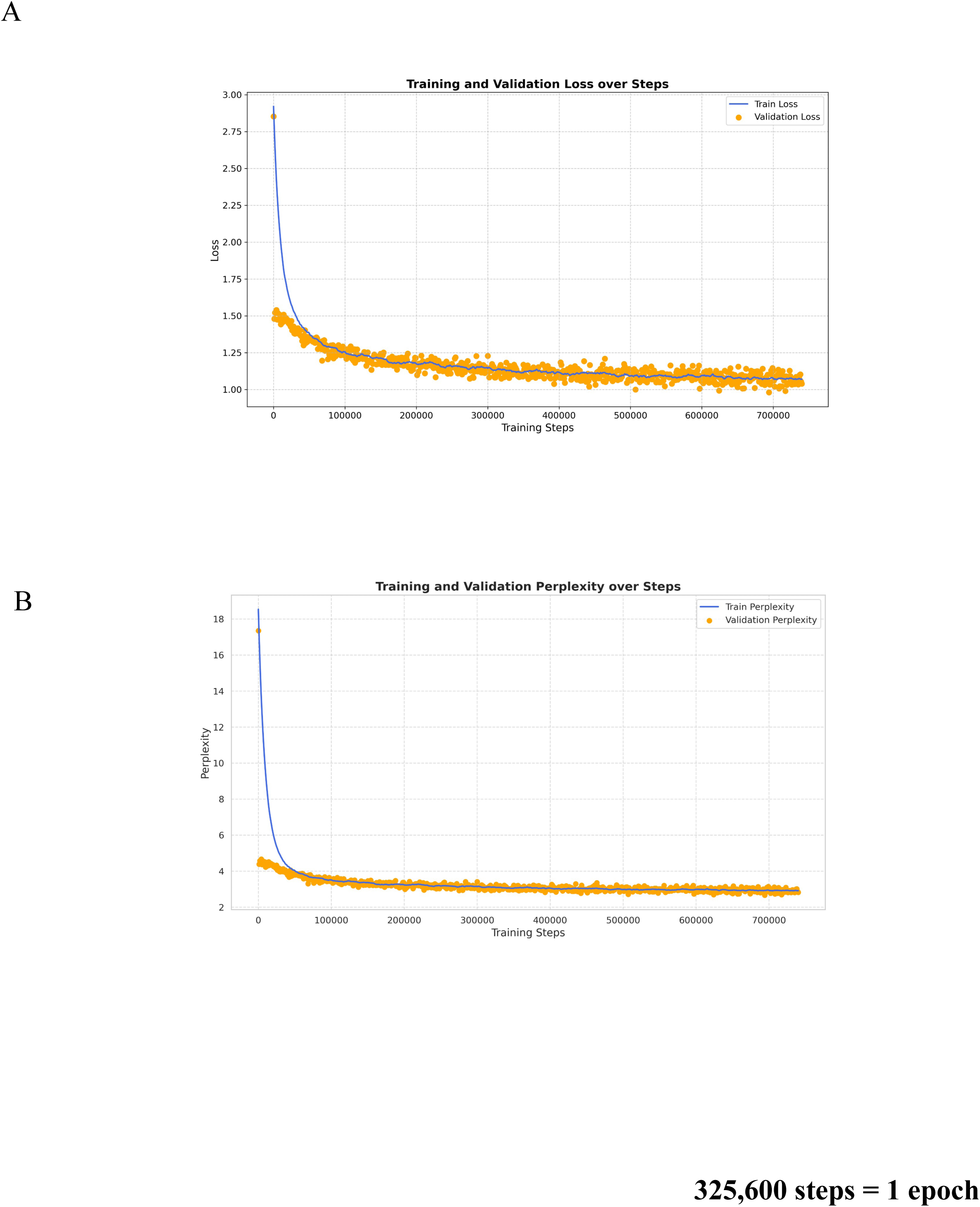
Loss and perplexity during pre-training of mRNA-GPT-eukaryote.

**Figure S3.**
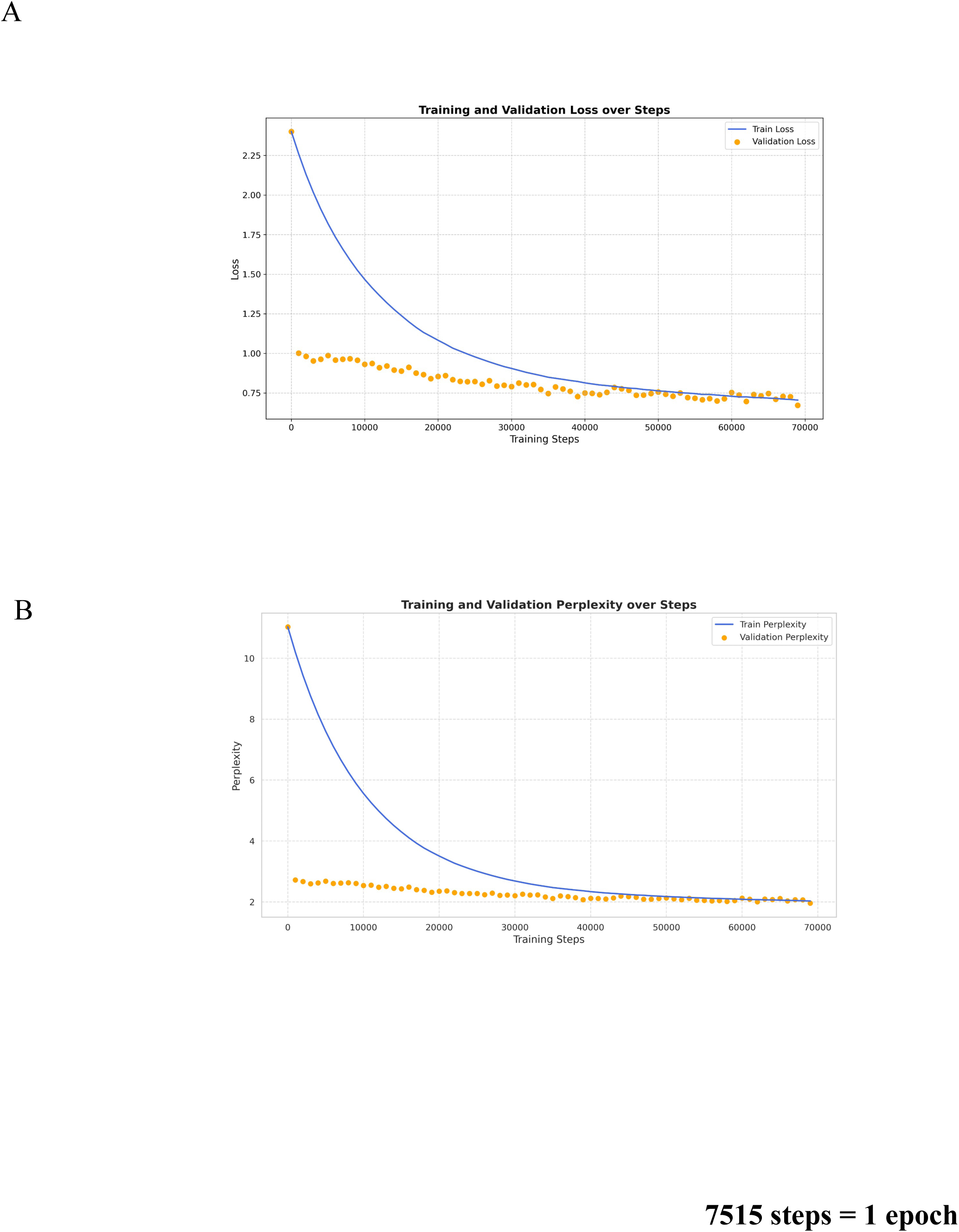
Loss and perplexity during pre-training of mRNA-GPT-archaea.

**Figure S4.**
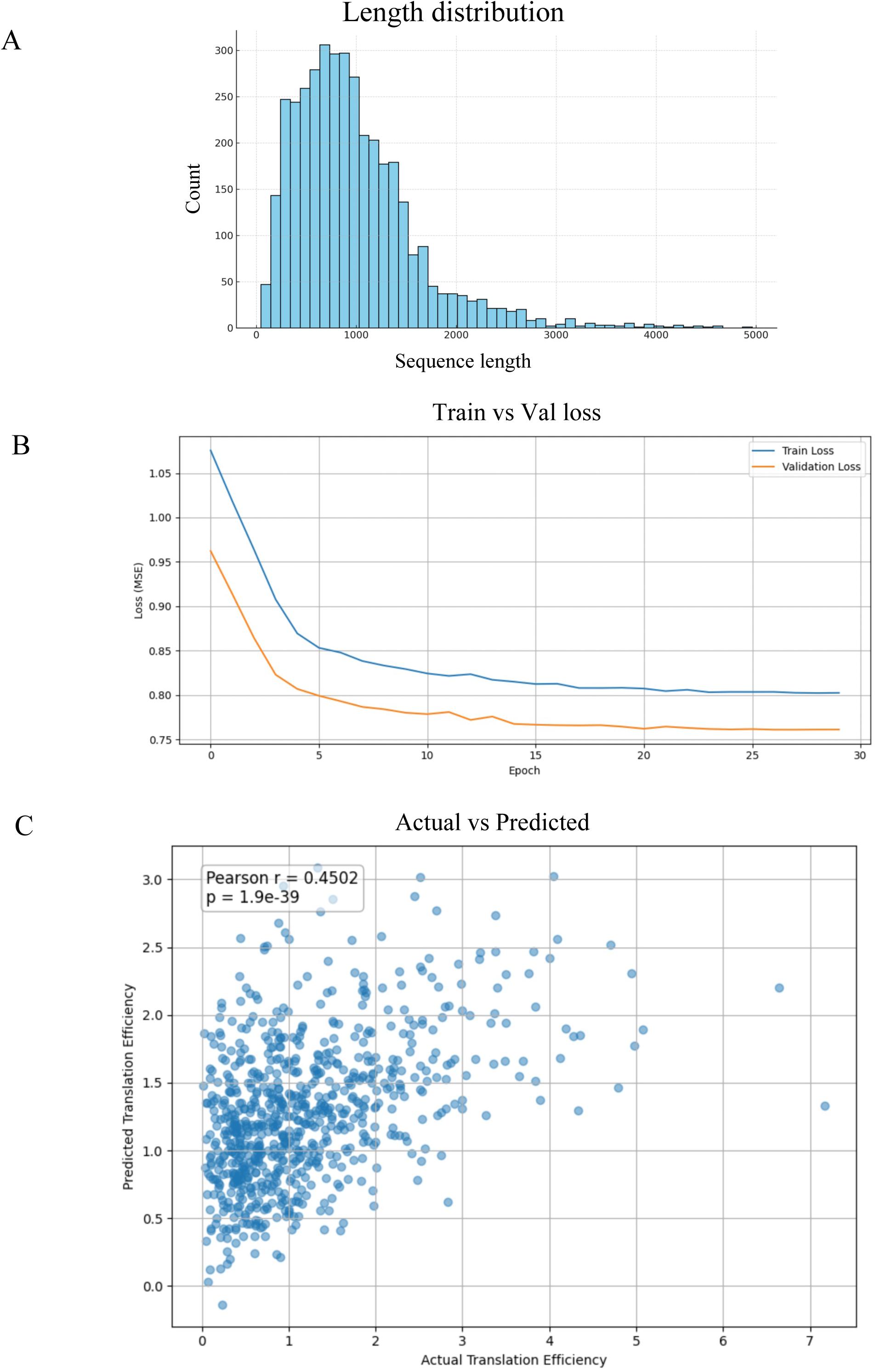
Training and performance of mRNA translation efficiency prediction model.

**Figure S5.**
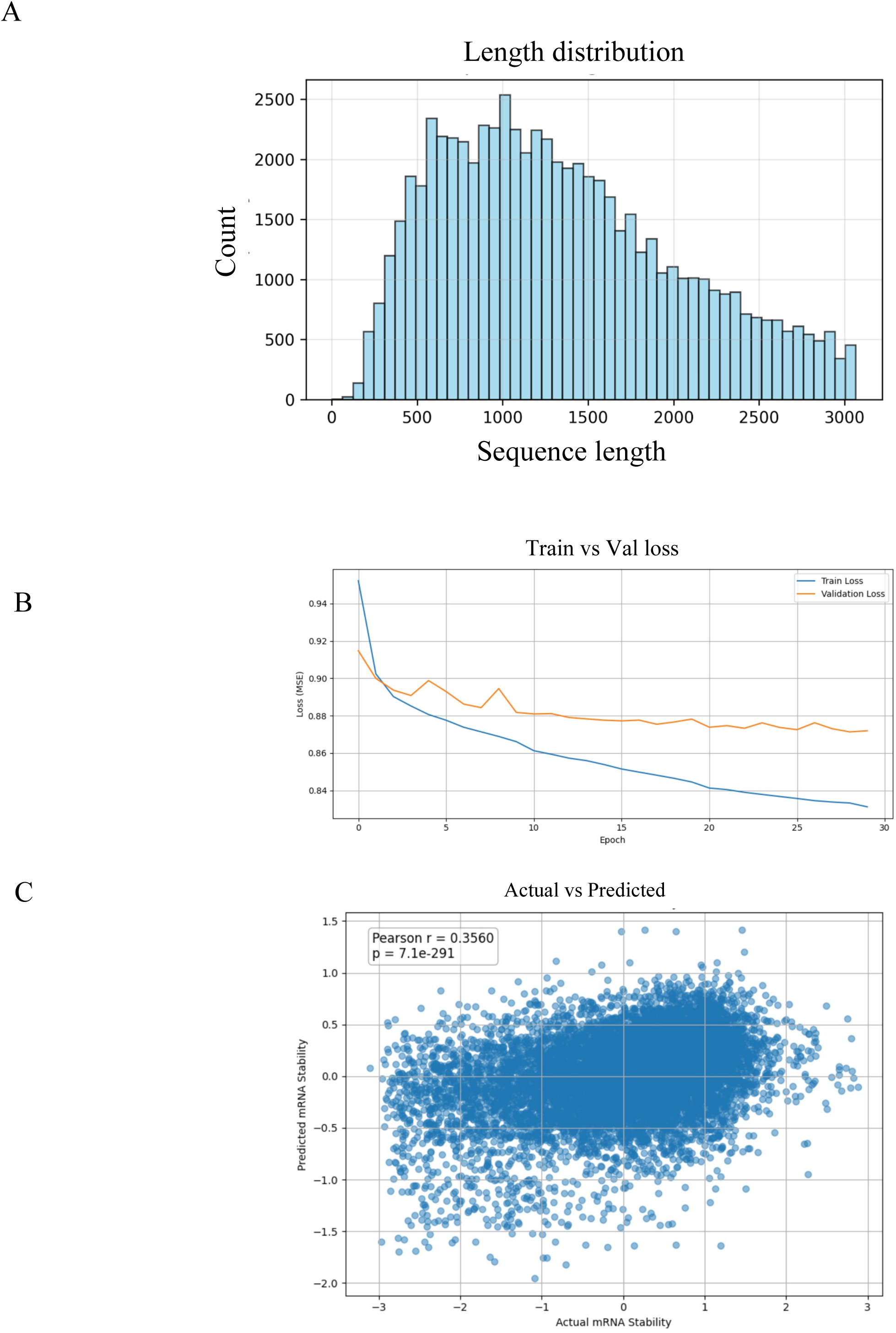
Training and performance of mRNA stability prediction model.

**Figure S6.**
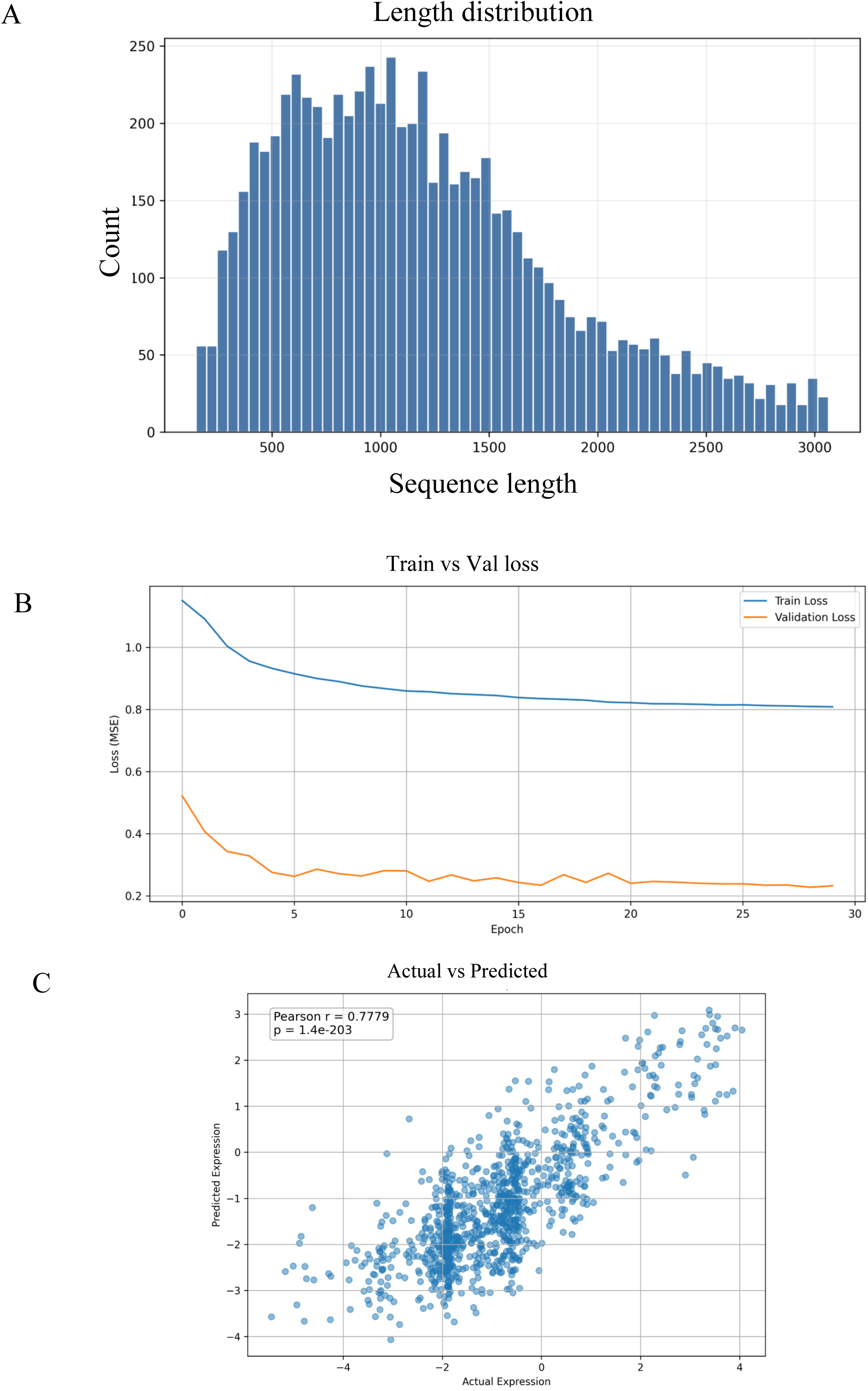
Training and performance of mRNA expression prediction model.

**Figure S7.**
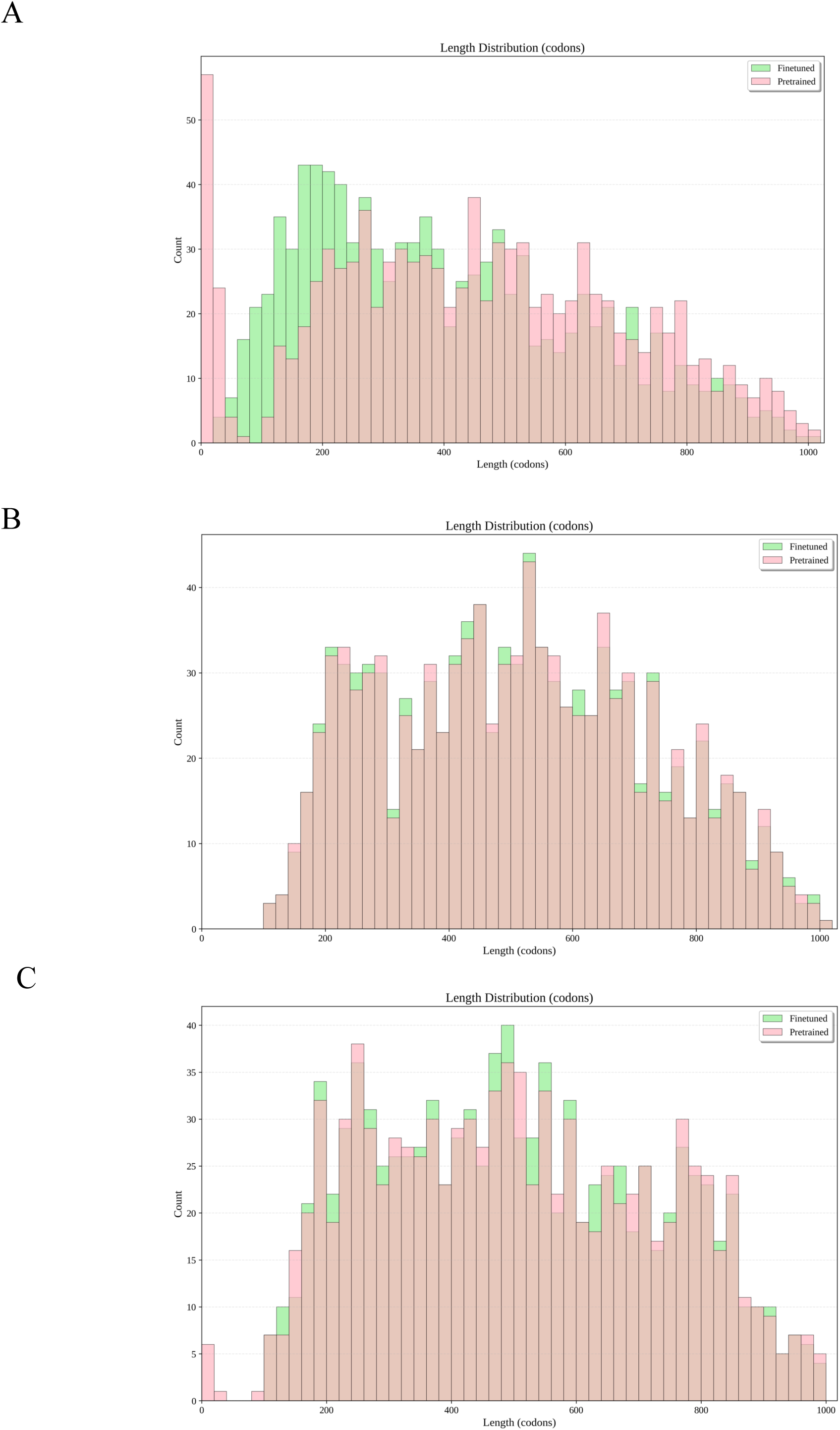
Length distribution of generated mRNA sequence from pretrained and finetuned model respectively (A) Bacteria, (B) Archaea, (C) Eukaryote.

